# MELK controls tumor metabolism to promote resistance to melanoma therapy

**DOI:** 10.64898/2025.12.31.697178

**Authors:** Ana Carolina B. Sant’Anna-Silva, Patricia Abbe, Sophie Pagnotta, Nivea D. Amoedo, Michael D. Torno, Fanny Bonnard, Thomas Botton, Frédéric Larbret, Cyril Ronco, Henri Montaudié, Issam Ben-Sahra, Caroline Robert, Thierry Passeron, Rodrigue Rossignol, Michaël Cerezo, Stéphane Rocchi

## Abstract

Melanoma is the most aggressive form of skin cancer, and despite major advances in targeted and immune therapies, durable responses remain limited due to the emergence of resistance mechanisms. Metabolic reprogramming has emerged as a key driver of therapy resistance, allowing tumor cells to adapt to environmental and therapeutic pressures. Here, we identify the maternal embryonic leucine zipper kinase (MELK) as a critical mediator of resistance in melanoma. We demonstrate that MELK stimulates intracellular accumulation of amino acids, leading to activation of mTORC1 signaling and enhanced mitochondrial metabolism and biogenesis. This metabolic shift promotes resistance to immune checkpoint blockade. Importantly, preclinical MELK inhibition sensitizes resistant melanoma cells to immunotherapy, revealing a potential combinatorial strategy to overcome resistance. Our findings establish MELK as a central regulator of metabolic adaptation and therapeutic resistance in melanoma, highlighting its potential as a promising therapeutic target.

## INTRODUCTION

Melanoma is the most aggressive type of skin cancer and represents a major concern for public health. Even though significant progress has been made in past 20 years with the development of new combinatory therapies, cancer cells develop resistance mechanisms to survive and overcome treatment. In the case of melanoma, approximately 50 % of the cases involve mutation in BRAF^V600E/K^, causing to constitutive activation of the MAPK/ERK (RAS-RAF-MEK-ERK) cascade. Currently, BRAF-mutated patients are treated with targeted therapy including combinations of BRAF and MEK inhibitors (aka BRAFi and MEKi). Alternatively, patients can be treated with immune checkpoint inhibitors (ICI), such as monoclonal antibodies against the programmed cell death protein 1, PD-1 (or its ligand PD-L1) and the cytotoxic T-lymphocyte-associated protein-4, CTLA-4, where both activate antitumor T-cell response. With the advent of targeted therapy and later the use of ICI, patient survival has significantly improved despite durable responses are achieved only in 30-50 % of cases.

Among the intrinsic mechanisms responsible for therapy resistance, metabolic reprogramming plays a major role within tumor adaptation to limiting resources^1^. Previous studies showed that BRAFi suppress glycolysis via inhibition of hexokinase II and glucose transporters in BRAF^V600E^ mutated melanomas, and this effect is reversed by acquisition of resistance^2^. Other studies have suggested that therapy-resistant cells rely on glutamine over glucose as the main carbon source as compared to melanocytes or non-resistant melanoma cells^3–5^ and the *in vitro* binding of anti-PD-L1 antibody can as well modulate glucose homeostasis^6^. All these aspects illustrate the correlation between tumor energy metabolism, anti-tumor immunity and acquisition of resistance to therapies. Studies on human melanomas biopsies also showed the existence of two bioenergetic classes of tumors, with a high or a low reliance on oxidative phosphorylation (OXPHOS)^7^. Moreover, high-OXPHOS melanoma that overexpress the fatty oxidation trifunctional enzyme HADHA have a different response to immunotherapy^8^. Still, the molecular determinants of the high-OXPHOS profile remain undetermined in many cancers.

The maternal embryonic leucine-zipper kinase (MELK) is an oncogene from the AMPK family overexpressed in multiple cancers, including melanoma^9^. MELK promotes mitosis of melanoma cells via the NF-κB pathway^9^, and its expression is correlated with poor prognosis^10^. Moreover, MELK expression has been linked to increased resistance to BRAFi^9^ and anti-inflammatory type macrophages polarization^11^, which are known to undergo a metabolic switch towards oxidative metabolism.

Due to the extensive role of MELK in tumorigenesis, several inhibitors have been developed attempting to target MELK, being demonstrated poorly selective^12^. In contrast, upon a drug screening, our group have developed a small-molecule that is highly selective for MELK inhibition^13^, and can be used as an alternative to target MELK in combination with current standard of care.

In this study, we demonstrate for first time that the kinase MELK enhances mTORC1 pathway signaling via stimulation of amino acids intracellular accumulation, which in turn leads to increase in mitochondrial energy metabolism, and ultimately driving resistance to immunotherapy in melanoma. Our preclinical results suggest that MELK is a promising target in combination with standard of care therapies to overcome resistance and improve melanoma patients’ response to treatment.

## RESULTS

### MELK protein is overexpressed in melanoma and correlates with aggressiveness

We first investigated MELK expression across GDC-TCGA and Genotype-Tissue Expression (GTEx) Project datasets including several cancer types comparing normal versus tumor tissues. Our analyses show that MELK has significantly higher expression in tumor versus normal tissues (**Fig. 1a**). Analysis of mRNA sequencing data from melanoma patients confirmed that MELK expression is increased in melanoma tissues compared to normal skin and MELK mRNA content is increased upon melanoma transformation (**Fig. 1b**). Further, patients’ samples stratification according to the most frequent melanoma mutations (in BRAF, NF1 and RAS genes) from the SKCM (TCGA) dataset revealed that MELK expression is significantly increased in tumor samples compared to normal tissues independent of their mutational status (**Fig. 1c**). Further, we explored MELK expression in a single-cell mRNA sequencing dataset of residual persister cells isolated from patient-derived melanoma xenografts grown in the presence of BRAFi+MEKi^14^. The results identified that MELK level associates with a proliferative state in melanoma tumors (**Fig. 1d**). Therefore, we first performed a gain of function approach in melanoma cells that have low baseline MELK expression. MELK expression was measured in melanoma cell lines with various mutational status (**Sup. Fig. 1a**) and MELK overexpression was performed in cells that presented with low levels of endogenous MELK, such as SK-MEL-10 (NRAS^G61R^) and YUMM 1.1 (BRAF^V600E^) (**Fig. 1e**). Upon MELK overexpression, cells presented increased proliferation rates (**Fig. 1f**). Similar findings were observed in cells that have high MELK baseline levels, such as A375 melanoma cells (**Sup. Fig. 1b,c**). MELK overexpression also led to an increase in tumorigenic properties such as clonogenicity, anchorage-independent colony formation and spheroid formation (**Fig. 1g-i**). Cell with high MELK expression also kept growing after long exposure (up to 72 h) to serum starvation (**Sup. Fig. 1c**). We then injected human melanoma cells overexpressing MELK or controls in nude mice and observed a significant increase in tumor volume (**Fig. 1j**) and *ex vivo* area in MELK cells (**Fig. 1k**). In contrast, targeting MELK in MELK high cells rescued tumor growth down to control levels (**Sup. Fig. 2a-c**). Our data indicate that MELK stimulates proliferation, tumor formation properties and malignancy *in vitro* and *in vivo*.

**Figure 1:**
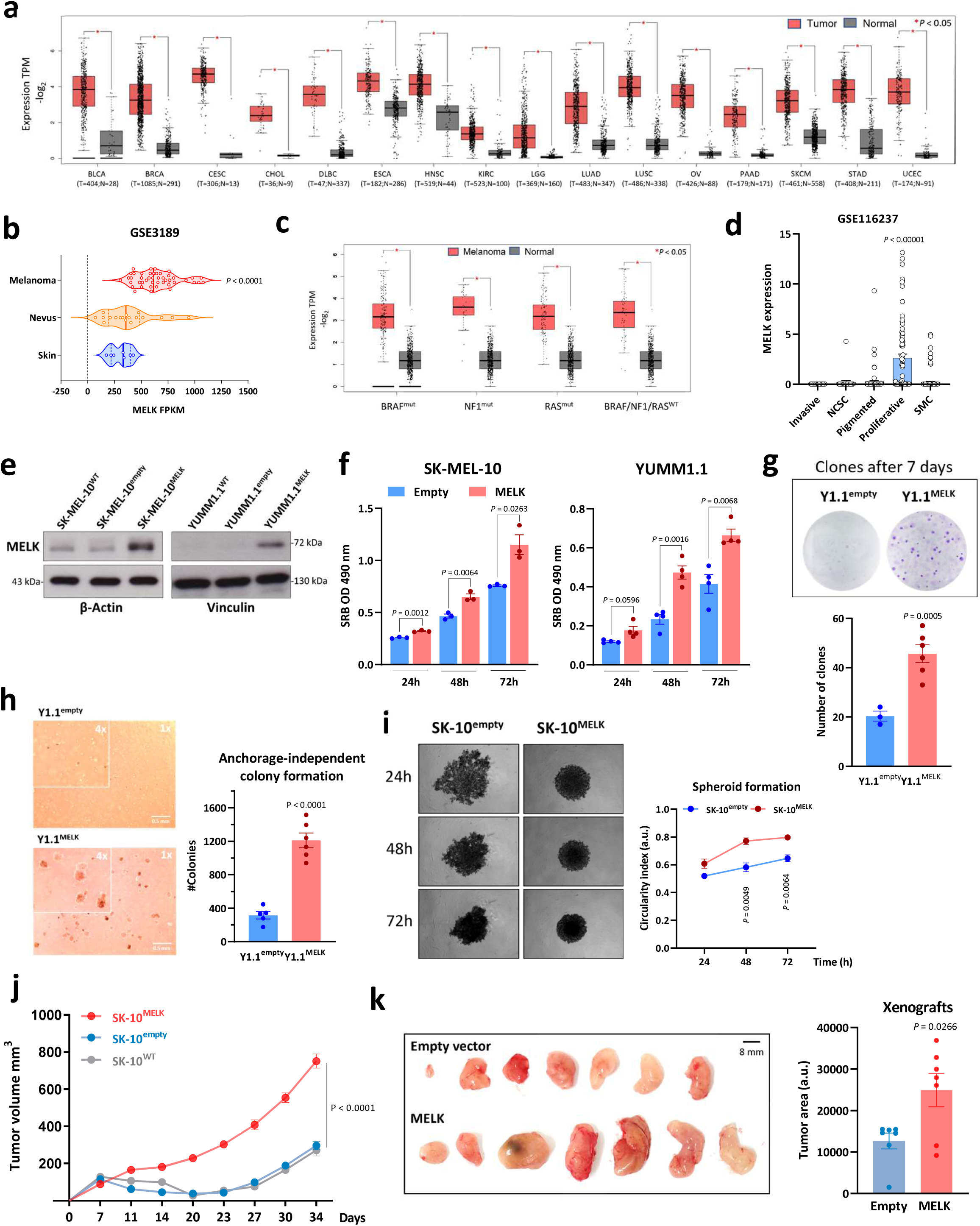
MELK is overexpressed in melanoma and its expression is correlated with aggressiveness. **(a)** MELK expression across several cancer types from TCGA datasets comparing tumor versus normal tissues; **(b)** Analysis of MELK mRNA expression using GSE3189 dataset between normal skin versus nevus or melanoma; **(c)** TCGA data base analysis using SKCM cohort stratifying patients wild-type or with BRAF, NF1 or RAS mutations and comparing MELK expression between melanoma and normal skin tissues; **(d)** Analysis of MELK mRNA expression using GSE116237 dataset in different persister cell states; **(e)** MELK overexpression in human SK-MEL-10 and murine YUMM1.1 melanoma cells; **(f)** Time course proliferation colorimetric assay over 72 h using sulforhodamine B in SK-MEL-10 and YUMM1.1 cells; **(g)** Clonogenic assay after 7 days incubation at basal culture conditions comparing Y1.1 cells expression empty vector or MELK; **(h)** Anchorage-independent colony formation assay; **(i)** Microspheroids formation over 72h incubation; **(j)** Immunodeficient BALB/c nude mice were inoculated subcutaneously with 5 × 10^6^ SK-MEL-10 LentiMELK, LentiControl (Empty vector) or WT cells. Tumor growth curves were determined by measuring the tumor volume; **(k)** *Ex vivo* tumor area measurements of the xenografts model. Bars indicate ± SEM; N ≥ 3; *P* values and biological replicates are indicated in the graphs.

**Figure 2:**
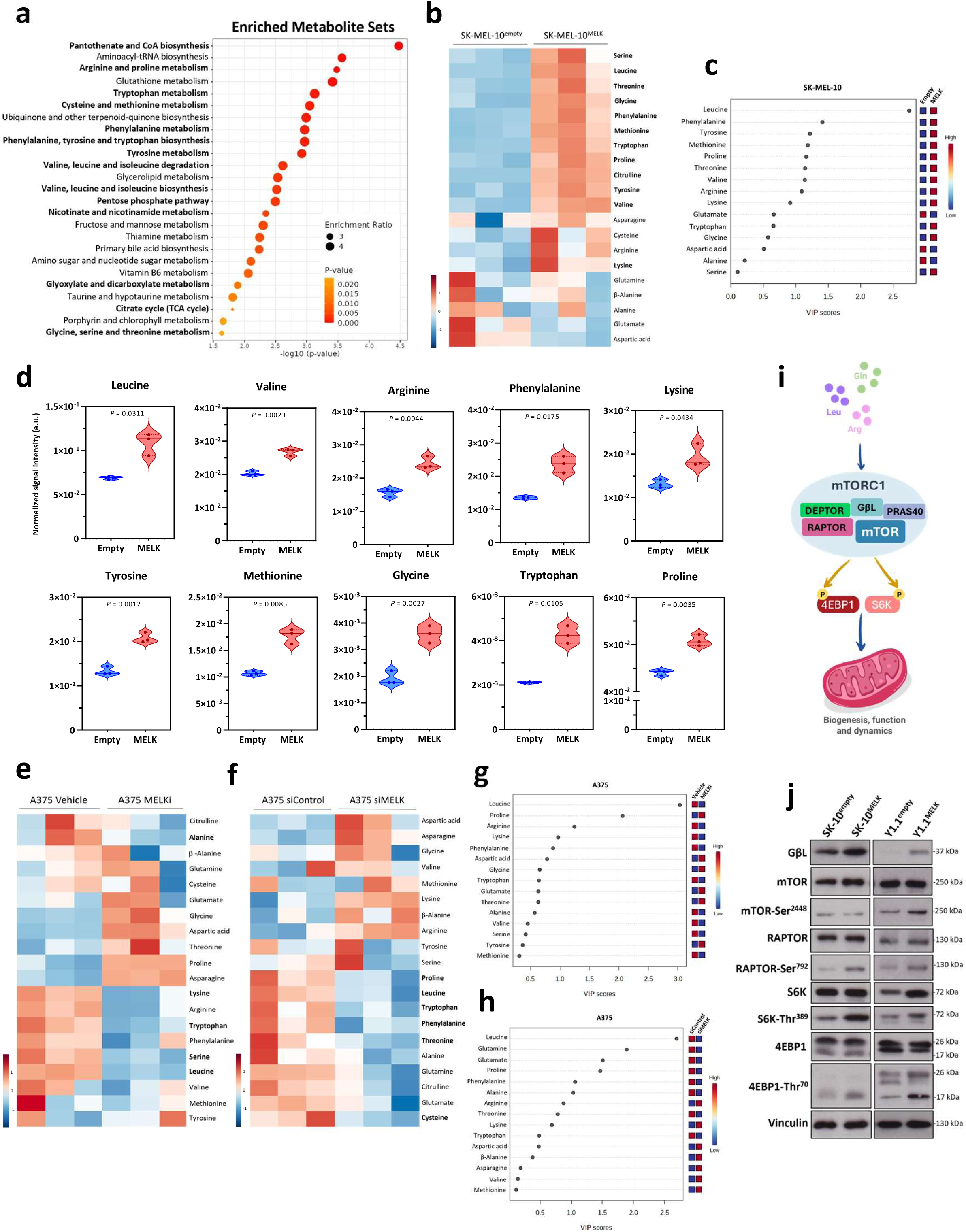
MELK expression stimulates amino acids accumulation and activates mTORC1 pathway. Untargeted metabolomics analysis from SK-MEL-10 expressing empty vector or MELK showing **(a)** Enriched metabolite pathways, **(b)** Heatmap showing amino acids accumulation, and **(c)** Variable Importance in Projection (VIP) score scale displaying the contribution of each amino acid to group discrimination; **(d)** Significant amino acids levels comparing SK-MEL-10 empty vector versus MELK; **(e)** Heatmap comparing A375 melanoma cells treated for 24h with 10 µM MELKi (CRO15) or vehicle; **(f)** Heatmap comparing A375 treated with 50 nM siRNA targeting scramble or MELK mRNA for 48h; **(g)** VIP scores from A375 cells treated 10 µM MELKi (CRO15) or vehicle; **(h)** VIP scores from A375 cells treated with 50 nM siRNA targeting scramble or MELK mRNA; **(i)** Illustration of amino acids activation of mTORC1 pathway that leads to downstream targets phosphorylation of 4EBP1 and S6K, promoting mitochondrial biogenesis, function and dynamics; **(j)** Immunoblots showing activation of mTORC1 pathway upon overexpression of MELK in SK-10 and Y1.1 melanoma cells. Violin plots indicate ± SEM; N ≥ 3; *P* values and biological replicates are indicated in the graphs.

### MELK stimulates amino acids accumulation and mTORC1 activation

The role of MELK in cancer metabolism remains undetermined, so we performed a series of unsupervised metabolomics and bioenergetic experiments. Steady-state metabolomics analysis showed that upon MELK overexpression, there was an enrichment of amino acids metabolism-related pathways, and most amino acids were accumulated intracellularly upon MELK overexpression comparing to control cells (**Fig. 2a,b**). VIP scores highlighted that the most important amino acid contributing to cluster separation between the samples was the brain-chain amino acids (BCAAs) leucine (**Fig. 2c**). However, several other amino acids presented increased levels in MELK high cells (**Fig. 2d**), in line with MELK chemical inhibition (MELKi) or siRNA against MELK, which led to decrease in amino acids concentrations (**Fig. 2e-h**). Since amino acids (especially BCAAs) feed mTOR enzymatic complexes, and mitochondrial metabolism and biogenesis can be regulated downstream mTORC1 (**Fig. 2i**), we analysed the mTORC1 pathway as well as mitochondrial physiology. We found that the main components of mTORC1 complex are upregulated in melanoma cells that overexpress MELK such as GβL, phospho-mTOR (S2448) and phospho-RAPTOR (S792), as well as their downstream targets phospho-S6K (T389) and phospho-4EBP1 (T70) (**Fig. 2j**), indicating that MELK activates mTORC1 complex. These observations show that MELK promotes accumulation of amino acids, especially BCAAs, which in turn activate mTORC1 signalling pathway.

### MELK expression stimulates mitochondrial function

To deeper investigate the metabolic reprogramming associated with MELK expression, we first analysed mitochondrial content using electron microscopy. The overexpression of MELK protein increased mitochondrial density as determined by organelle area in the cytosol (**Fig. 3a,b**). Mitochondrial function was also stimulated as shown by increase in basal mitochondrial respiration (**Fig. 3c**), Complex I activity (**Fig. 3d**), total dehydrogenase activity (**Fig. 3e**) and citrate synthase activity (**Fig. 3f**). An increase in mtDNA content was also determined in tumor xenografts extracted from mice transplanted with MELK overexpressing cells or empty vector (**Fig. 3g**). Mitochondrial respiratory complexes capacities measured *ex vivo* by high-resolution respirometry also showed an increase in the maximal capacity of Complexes I and Complex II-dependent respiratory systems (NADH-pathway and succinate-pathway respectively) (**Fig. 3h,i**). These findings evidence the role of MELK in the control of mitochondrial biogenesis and function. Considering that mTORC1 stimulates mitochondrial biogenesis, we checked the correlation between MELK expression and mitochondria-controlling transcription factors using a cutaneous melanoma TCGA cohort with 470 patients’ samples. This analysis showed that MELK mRNA expression positively correlates with both NRF1 and TFAM expressions in melanoma tumors, as well as SIRT1 (**Sup. Fig. 3a-c**) but not PGC1α (PPARGC1A gene) (**Sup. Fig. 3d**). We confirmed *in vitro* that upon MELK overexpression there is an increase NRF1 and TFAM, as well as SIRT1 protein levels (**Fig. 3j**). Our results demonstrate that MELK cells stimulate mitochondrial metabolism, enhancing mitochondrial biogenesis and function.

**Figure 3:**
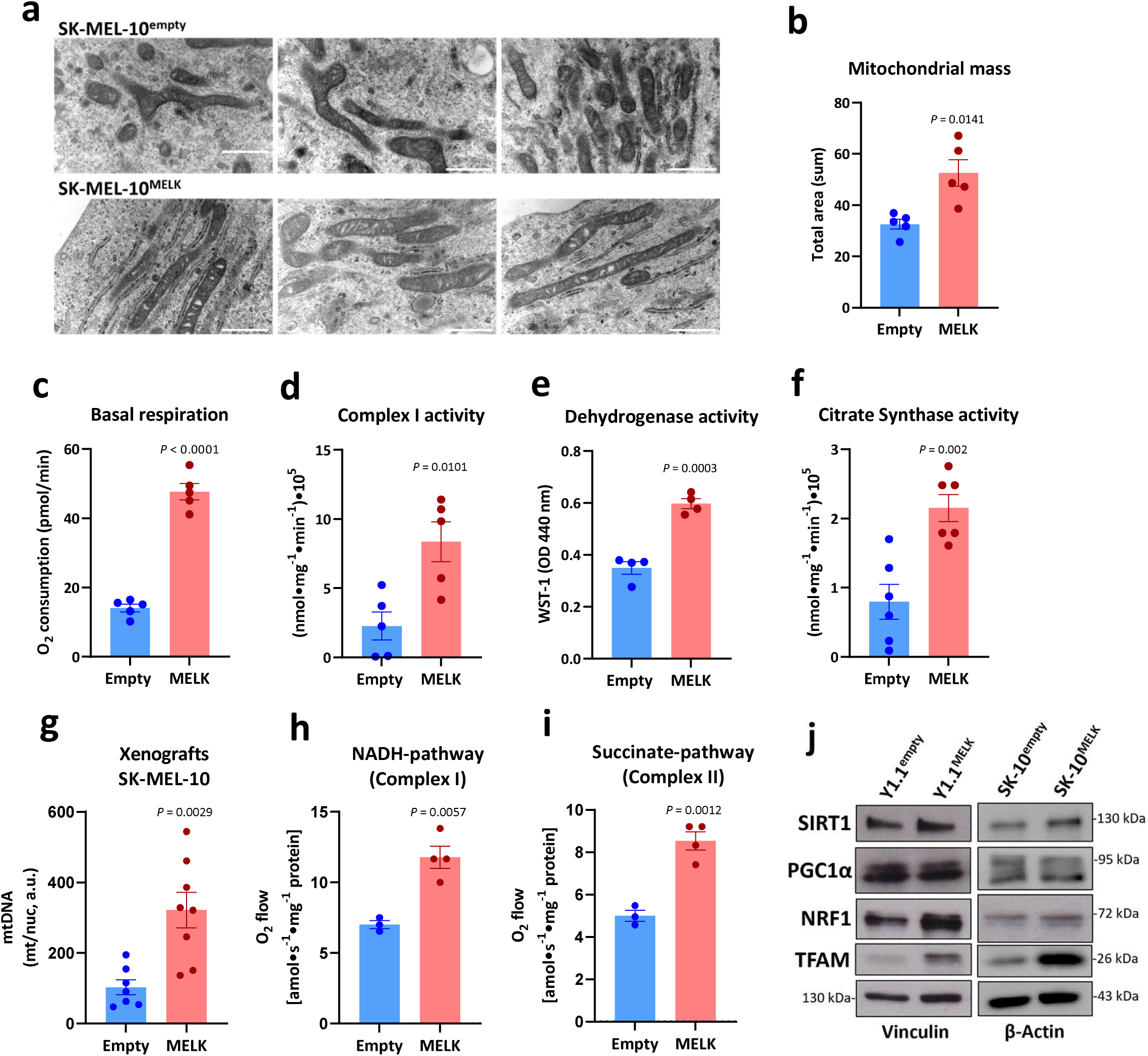
MELK activates mitochondrial function. **(a)** Electron Microscopy of Transmission showing mitochondrial network; **(b)** Quantification of mitochondrial mass by total area; **(c)** Oxygen consumption corresponding to basal respiration of SK-10 cells cultivated in full cell culture medium (pmol/min); **(d)** Complex I activity (nmol*min^−1^*min^−1^); **(e)** Dehydrogenase activity measured with WST-1 assay; **(f)** Citrate synthase activity (nmol*min^−1^*min^−1^); **(g)** mRNA expression ratio between mitochondrial and nuclear genomes corresponding to relative mtDNA quantification in total mRNA extracted from tumor xenografts inoculated with SK-MEL-10 cells; Mitochondrial **(h)** NADH and **(i)** Succinate pathways capacities from tumor xenografts homogenates (nmol*min^−1^*mg^−1^ protein); **(j)** Immunoblotting showing mitochondrial biogenesis proteins in YUMM1.1 Empty vector or MELK and SK-MEL-10 Empty vector or MELK overexpression. Bars indicate ± SEM; N ≥ 3; *P* values and biological replicates are indicated in the graphs.

### Targeting MELK leads to decrease in mitochondrial metabolism

To check whether by targeting MELK we could potentially revert enhanced metabolic phenotype, we have used the MELK inhibitor CRO15^13^. Our results showed that MELK inhibition leads to a decrease of mitochondrial complexes subunits expression in SK-10^MELK^ cells compared to empty controls and that our inhibitor is specific for cells with high MELK levels overtime (**Fig. 4a**). Experiments were also performed with another MELKi, MELKT1^15^ (data not shown). MELK inhibition effect on mitochondrial subunits expression was confirmed in A375 cells with high basal level of MELK expression, using metformin as a mitochondrial inhibitor positive control (**Fig. 4b**), as well as in MELK knockout melanoma cells (**Fig. 4c**). MELK chemical inhibition also promoted mitochondrial cristae partial loss and matrix swelling, which is an indication of mitochondrial stress in MELK high cells (**Fig. 4d**). Basal respiration was also diminished in MELK knockout A375 cells (**Fig. 4e**). Furthermore, targeting MELK high tumors led to the inhibition of mitochondrial substrate oxidation capacity through Complexes I and II and the supercomplex I+II in tumor xenografts homogenates (**Fig. 4f-h**). Analysis of glucose metabolism in MELK high cells showed a reduction of glycolysis intermediates, suggesting a decrease in this pathway (**Sup. Fig. 4a**). Lactate dehydrogenase activity (LDH) was not altered by MELK overexpression (**Sup. Fig. 4b**) and MELK high cells increased extracellular lactate release upon mitochondrial inhibition (**Sup. Fig. 4c**), suggesting a higher dependency on OXPHOS. Taken together, our results demonstrate that MELK expression induces a metabolic shift towards mitochondrial metabolism, specifically stimulating a metabolic rewiring program that favours oxidative over glycolytic function.

**Figure 4:**
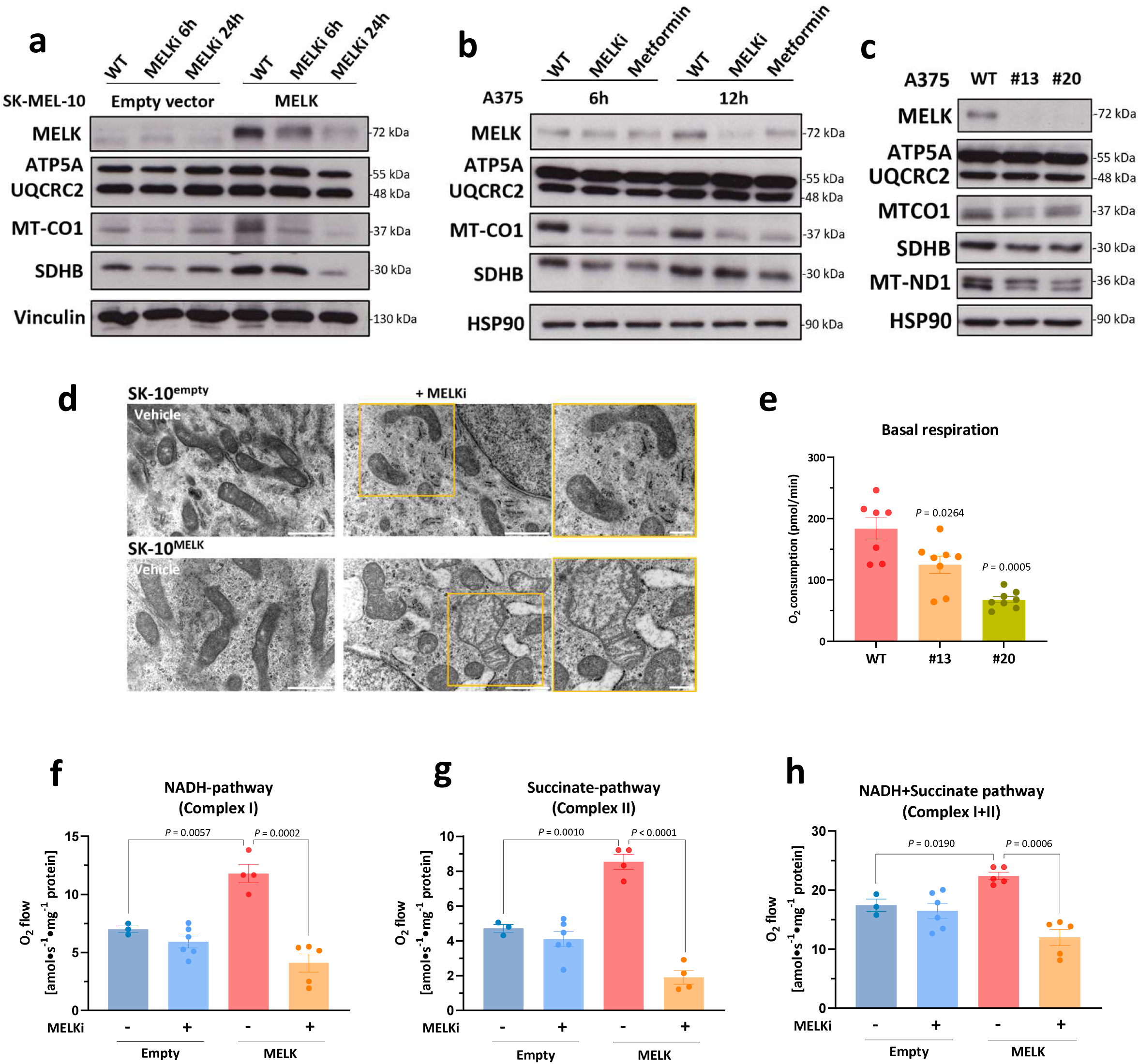
MELK targeting inhibits mitochondrial function. **(a)** Immunoblot displaying respiratory complexes expression of SK-MEL-10 cells expressing empty vector or MELK plasmids treated or not with MELKi (CRO15) for 6h or 24h; **(b)** Immunoblot displaying respiratory complexes expression of A375 cells treated for 6h or 12h with 10 µM MELKi (CRO15) or metformin; **(c)** Protein expression of mitochondrial respiratory complexes of A375 melanoma cells CRISPR-Cas9 knocked-out for MELK with two different clones (#13 and #20) or wild-type (WT); **(d)** Electron Microscopy of Transmission showing mitochondria shape in SK-MEL-10 Empty vector or MELK treated with vehicle or MELKi (CRO15) for 24h; **(e)** Oxygen consumption of A375 CRISPR-Cas9 knock-out cells; Mitochondrial **(f)** NADH, **(g)** Succinate, and **(h)** NADH+Succinate pathways capacities from tumor xenografts homogenates (nmol*min^−1^*mg^−1^ protein). Bars indicate ± SEM; N ≥ 3; *P* values and biological replicates are indicated in the graphs.

### MELK expression is associated with disease progression, linked to lower immune infiltration and resistance to melanoma therapy

Mitochondrial biogenesis and TFAM expression have been previously associated with decreased anti-tumor immune response and resistance to therapies in melanoma^16,17^. Public dataset analysis revealed an increase in MELK mRNA levels in melanoma patients with disease progression compared to those who responded to treatment (**Fig. 5a**), as well as in patients who have received immunotherapy (**Fig. 5b**). Moreover, patients with high MELK expression had lower survival curves after ICI treatment and MELK expression has been associated with higher immune resistance and lower immune infiltration in different cohorts of patients (**Fig. 5c-e**). To understand whether the consequences of elevated MELK expression might lead to a phenotype of resistance, we proceeded by injecting YUMM1.1 melanoma murine cells overexpressing MELK or empty vector in immunocompetent mice and after tumor reached approximately 100 mm^3^, we started treating the mice with ICI combination anti-PD1 + anti-CTLA-4. MELK tumors grew much faster than controls and did not respond to ICI, unlike control tumors (**Fig. 5f,g**). These results suggest that MELK promotes resistance to immunotherapy in melanoma.

**Figure 5:**
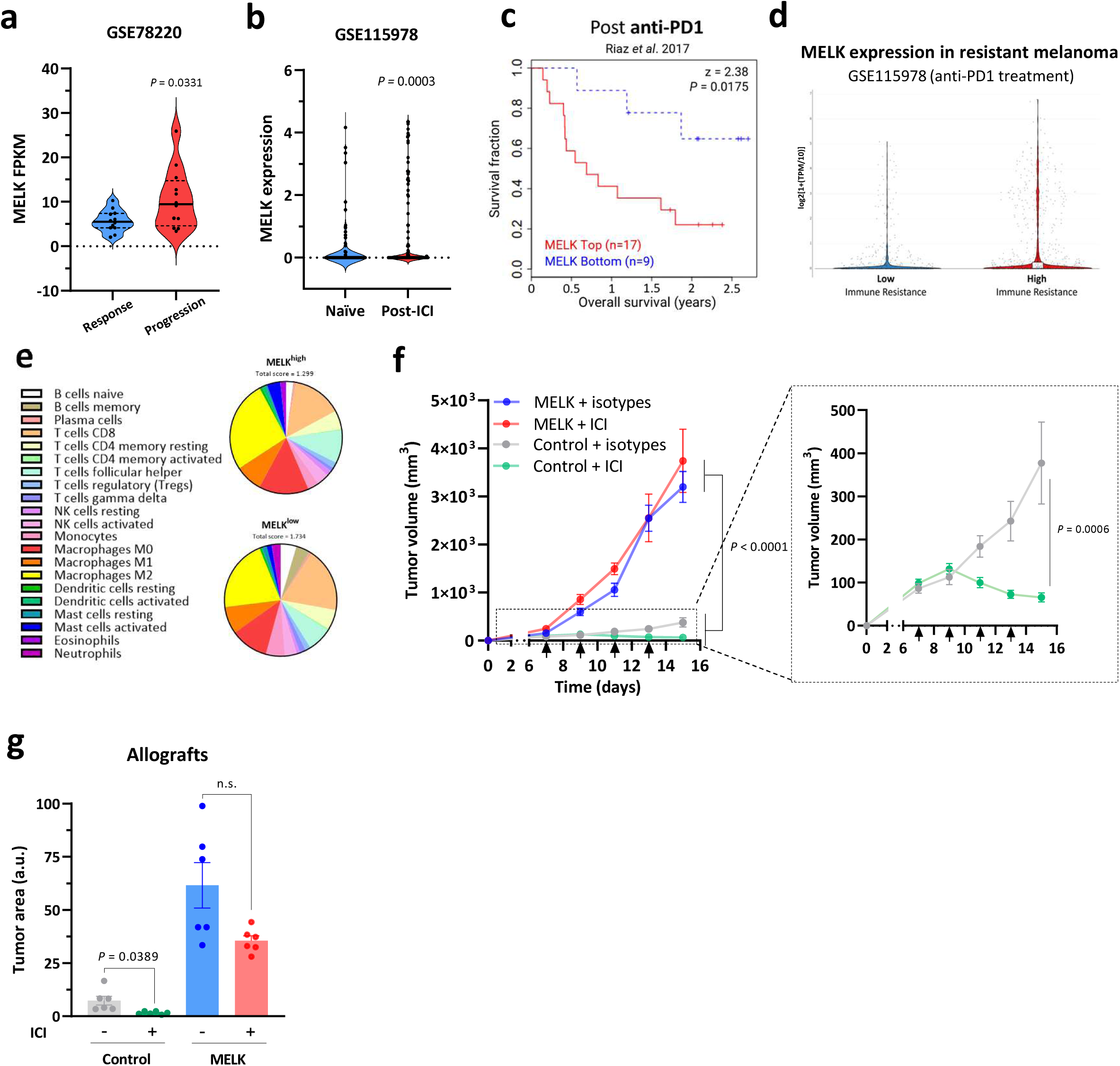
MELK expression promotes resistance to immune checkpoint blockade. **(a)** MELK mRNA expression across samples of patients that responded or not (progression disease) after immunotherapy using GSE78220 dataset; **(b)** MELK mRNA expression comparing naïve and post immune checkpoint inhibitor (post-ICI) treatment using GSE115978 dataset; **(c)** Survival curves after anti-PD1 treatment stratifying for patients with MELK high (N = 17) or low expression (N = 9); **(d)** MELK expression in resistant melanoma samples after anti-PD1 treatment classified into low or high immune resistance; **(e)** *In silico* analysis using GSE78220 dataset comparing the immune cells infiltration prediction between MELK^low^ and MELK^high^ melanoma samples; **(f)** Immunocompetent C57Bl6/J mice were inoculated subcutaneously with 2 × 10^6^ YUMM 1.1 cells overexpressing empty vector (Control) or MELK. Mice were treated with anti-PD-1 + anti-CTLA-4 (100 µg/mouse/day) every 2 days. Tumor growth curves were followed by measuring the tumor volume (mm^3^); **(g)** *Ex vivo* tumor area measurements of the allografts model. Bars indicate ± SEM; N ≥ 3; *P* values and biological replicates are indicated in the graphs.

### Combination therapy with the targeting of MELK reverts MELK-associated phenotype

To further investigate the effects on immunotherapy response exerted by high MELK levels, we have inoculated MELK-overexpressing murine melanoma cells in C57BL/6 immunocompetent mice in parallel with controls cells. After tumors had reached approximately 100 mm^3^ we treated the mice with MELKi or vehicle. Results show that MELKi selectively targeted MELK-overexpressed cells in contrast with controls and when used as a single treatment, led to decrease in tumor volumes compared to vehicle-treated tumors (**Fig. 6a**). We also observed a that MELKi treatment response is potentialized by ICI treatment, promoting a decrease in tumor volumes and areas (**Fig. 6b,c**), contrasting with no effect observed in empty vector tumor, that as expected, responded well to ICI at basal levels (**Fig. 6d,e**). Confirming the metabolic downstream effect of MELK inhibition, isolated tumors treated with MELKi+ICI presented lower mitochondrial DNA markers expression (**Fig. 6f**), suggesting lower mitochondrial function upon MELK inhibition *in vivo*. These results indicate that targeting MELK in combination with ICI might overcome MELK phenotype associated with mitochondrial metabolic shift, ultimately contributing to bypass resistance in melanoma treatment and presenting a new potential therapy to combat therapy resistance in MELK high melanomas.

**Figure 6:**
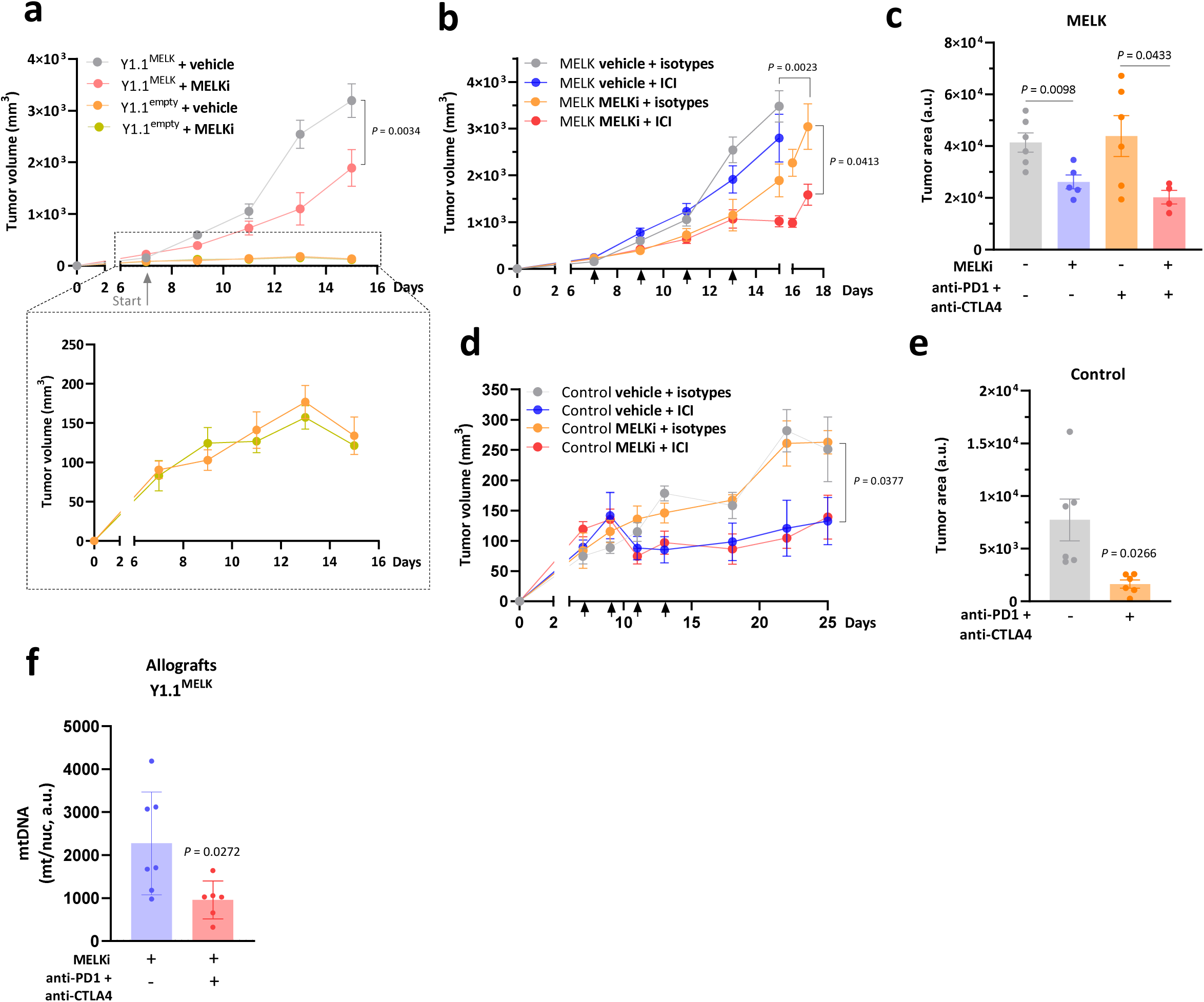
Targeting MELK in combination with immunotherapy resensitize MELK resistant tumors. Immunocompetent C57Bl6/J mice were inoculated subcutaneously with 2 × 10^6^ YUMM 1.1 cells overexpressing empty vector (Control) or MELK. **(a)** Mice were treated with MELK inhibitor CRO15 (35 mg/kg/day) daily or vehicle; Mice were treated with MELK inhibitor CRO15 (35 mg/kg/day) daily or vehicle plus anti-PD-1 + anti-CTLA-4 (100 µg/mouse/day) or isotypes every 2 days for **(b)** MELK tumors or **(d)** Control tumors. Tumor growth curves were followed by measuring the tumor volume (mm^3^); *Ex vivo* tumor area measurements of the allografts model in **(c)** MELK tumors or **(e)** Control tumors; **(f)** mRNA expression ratio between mitochondrial and nuclear genomes corresponding to relative mtDNA quantification in total mRNA extracted from tumor allografts inoculated with YUMM1.1 cells overexpressing MELK. Bars indicate ± SEM; N ≥ 3; *P* values and biological replicates are indicated in the graphs.

## DISCUSSION

Despite remarkable advances in melanoma treatment with targeted therapies and immune checkpoint blockade (ICB), durable responses remain elusive for a substantial fraction of patients, largely due to the emergence of resistance mechanisms^18,19^. Here, we identify MELK as a key regulator of therapy resistance, acting through increase in intracellular amino acids to sustain mTORC1 signaling and promoting mitochondrial biogenesis and enhanced function. This metabolic adaptation enables melanoma cells to persist under the selective pressures imposed by ICB. Importantly, pharmacologic, or genetic targeting of MELK sensitizes resistant melanomas to therapy, establishing MELK as a potential biomarker of resistance.

MELK has historically been described as a mitotic kinase that supports proliferation and oncogenesis in multiple cancers, including melanoma, glioblastoma, and breast cancer^20–22^. Its prognostic value has been linked to aggressive disease and poor clinical outcome^23,24^. However, its role in metabolic adaptation has not yet been elucidated. Our data reveal that MELK drives a profound metabolic shift toward mitochondrial dependency, enhancing mitochondrial mass, electron transfer system, and OXPHOS capacity. This MELK-driven reprogramming may account for the prior observations that therapy-resistant melanomas transition away from glycolysis and towards mitochondrial metabolism to support bioenergetic and biosynthetic needs^4,7,25^.

mTORC1 is a master regulator of metabolic homeostasis, linking amino acid availability to mitochondrial biogenesis and oxidative metabolism^26,27^. By sustaining mTORC1 activity, MELK promotes mitochondrial expansion and transcriptional activation of TFAM and NRF1, reinforcing OXPHOS dependency. These data establish MELK as a previously unrecognized metabolic regulator in melanoma.

Therapy resistance in melanoma is multifactorial, involving both cell-intrinsic mechanisms and microenvironmental influences^28,29^. Our findings place MELK at the intersection of these processes. MELK^high^ tumors exhibited decreased tumor-infiltrating lymphocytes, particularly CD8+ T cells, and failed to respond to ICI *in vivo*. These observations are in line with prior reports that metabolic rewiring toward OXPHOS supports an immune-suppressive tumor microenvironment and resistance to immunotherapy^30–32^. Thus, MELK promotes therapy resistance by coupling metabolic reprogramming with immune evasion, a dual mechanism that helps explaining its strong association with poor patient survival and disease progression across clinical datasets.

Metabolic rewiring has long been recognized as a hallmark of cancer. However, its targeting has not yet been successfully translated into clinical settings. Our work identifies a novel regulator of mTORC1 signaling and mitochondrial metabolism, offering a new potential therapeutic target to inhibit tumor cell metabolic plasticity and improve treatment efficacy. Moreover, our findings position MELK as a key driver of metabolism-driven resistance, suggesting its potential utility as an early biomarker to stratify patients at risk of developing resistance.

In conclusion, our work identifies MELK as a central regulator in tumor metabolic reprogramming and immunotherapy resistance in melanoma. By sustaining mTORC1 signaling and mitochondrial biogenesis, MELK enable tumor persistence under therapy. MELK inhibition reverses these adaptations and restores therapy sensitivity, revealing a promising therapeutic strategy to overcome resistance. More broadly, the MELK–mTORC1 axis represents a fundamental mechanism of tumor metabolic plasticity and a novel target for intervention in cancer.

## METHODS

### Cell lines and culture

The human melanoma cell lines A375, SK-MEL-28, MEL-501, MEWO, A2058, MEL-624 and SK-MEL-10 used in this study were purchased from the American Type Culture Collection. The human melanoma cell lines M117 and M6, and the murine melanoma cells 1007, 1014 and B16F10 from were a gift from Sophie Tartare-Deckert (INSERM U1065, France). The murine melanoma cell line YUMM 1.1 was a gift from Mehdi Khaled (Gustave Roussy, France). Melanoma cells were maintained at 37 °C and 5 % CO_2_ in a humidified atmosphere and grown in RPMI 1640 or DMEM growth medium supplemented with 10 % FBS. All cell lines were regularly tested for mycoplasma-free. Cells were treated with MELK inhibitors MELKT1^15^ (MedChemExpress) or CRO15^13^ at concentrations and time-points indicated in the figures.

### Lentiviral transductions for stable cell line generation

To overexpress MELK in human melanoma cells, we used pLX304-Blast-V5 vector expressing human MELK gene taken from the CCSB-Broad Lentiviral Expression Collection (Horizon Discovery). The plasmid pLX304 #25890 from Addgene was used as empty control. To overexpress MELK in murine melanoma cells, mCherry vector backbone were designed by Vector Builder (Chicago, USA) using as a template pLV[Exp]-EF1A>mCherry (VB900088-2479xcj) vector, that was also used as empty control.

Plasmids were transfected along with lentivirus packaging DNA using JetPRIME transfection Reagent Kit (Polyplus) into human embryonic kidney HEK 293T cells. After 6 h transfection, the medium was changed, and supernatants were collected after 48 h. After centrifugation, lentivirus production was mixed and frozen containing 8 µg/mL Polybrene (Sigma-Aldrich). Cells were infected with lentiviruses supernatants and selected either with antibiotics for ∼20 days (pLX304 vectors) or cells were sorted for mCherry expression (pLV-mCherry vectors).

### Animal studies

The experiments were performed in accordance with the Declaration of Helsinki and were approved by a local ethical committee CIEPAL (Comite Institutionnel d‘Ethique Pour l’Animal de Laboratoire-Azur). All mice used in this study were housed under pathogen-free conditions with food and water ad libitum. Animals were blindly allocated to experimental groups so that the groups had similar mean tumor volumes prior to treatment initiation.

For the xenografts’ model, female immune-deficient BALB/c nu/nu (nude) mice were obtained at 5 weeks of age from Envigo Laboratory (Gannat, France). At 6–7 weeks of age, the mice were injected subcutaneously into the flank with 5 × 10^6^ SK-MEL-10 LentiMELK, LentiControl (Empty vector) or WT cells with 50 % Matrigel® Matrix (Corning #354230). For MELKi tumor growth experiments, after tumors reached ∼100 mm^3^ (approx. 1 week later), animals received intraperitoneal injection with Labrafil (Control) or CRO15 (0.7 mg/mouse/day) dissolved in Labrafil.

For the allografts’ model, male immunocompetent C57Bl/6JOlaHsd mice were obtained at 5 weeks of age from Envigo. At 6–7 weeks of age, the mice were injected subcutaneously into the flank with 2 × 10^6^ YUMM 1.1 cells overexpressing empty vector (Control) or MELK. Mice were treated with anti-PD-1 + anti-CTLA-4 (100 µg/mouse/day) or isotypes controls every two days. For MELKi tumor growth experiments, mice treated with anti-PD-1 + anti-CTLA-4 or isotypes every two days were also treated with intraperitoneal injections of either Labrafil (Control) or CRO15 (0.7 mg/mouse/day) daily.

Tumor growth was monitored at minimum three-times a week in two dimensions using a digital calliper. The tumor volumes were calculated using the following ellipsoid volume formula: V = (L×W2)/2, where L is the length and W is the width.

### In silico data sets analyses

*In silico* analyses investigating MELK expression in tumor tissues versus normal and correlation among melanoma mutational statuses were performed using cBioPortal (https://www.cbioportal.org/ Accessed on 17.12.2025) with the TCGA GDC Cutaneous Melanoma cohort using 473 samples from 470 patients. MELK expression in melanoma malignant transformation, persister cell states and immune checkpoint resistance was done using the following the datasets: GSE3189^33^, GSE116237^14^, GSE78220^29^, GSE115978^34^.

### Protein extraction and immunoblotting

Cells were seeded in 6-well plates at approximately 0.25 × 10^6^ cells/well and harvested after 24 h incubation at 70–80 % confluency unless otherwise stated. Cells were washed with PBS and 80 µL of 1 × RIPA buffer (Sigma-Aldrich) with protease and phosphatase inhibitors (Merck) was added in each well. Cells were scraped-off, collected and frozen at −80 °C until further use. For protein extraction and immunoblotting, cell lysates were thawed, and proteins were quantified using Pierce BCA protein assay kit. LDS Sample Buffer (Thermo Fisher Scientific) was added, and sample volume was adjusted to equal protein concentration. Samples were heated at 95 °C for 5 min and 20–30 µg of protein extract per lane was loaded on a 7.5-12 % SDS-PAGE gel depending on the molecular weight of the blotted proteins. After running at 100 V for 1.5 h, the SDS-PAGE gels were transferred onto a polyvinylidene fluoride membrane (Millipore) at 100 V for 2 h. The membranes were then placed in Ponceau protein stain solution (Thermo Fisher), washed, blocked with BSA in TBS-Tween 1 % for 1 h and then incubated with primary antibodies diluted in with BSA in TBS-Tween 1 % overnight at 4 °C. The following primary antibodies were used for immunoblotting: anti-MELK (1:1000 Cell Signalling Technologies #2274S); anti-vinculin (1:1000 Sigma-Aldrich #V9131); anti-β-actin (1:1000 Santa Cruz Biotechnology #sc-47778); total OXPHOS Rodent WB Antibody Cocktail (1:1000 Abcam #ab110413, containing the antibodies anti-NDUFB8, anti-SDHB, anti-UQCRC2, anti-COX II and anti-ATP5 as a premixed solution); anti-GβL (1:1000 Cell Signalling Technologies #3274T); anti-mTOR (1:1000 Cell Signalling Technologies #2983S); anti-phospho-mTOR (1:1000 Cell Signalling Technologies #2971S); anti-RAPTOR (1:1000 Cell Signalling Technologies #48648); anti-phospho-RAPTOR (1:1000 Cell Signalling Technologies #89146); anti-p70 S6 kinase (1:1000 Cell Signalling Technologies #9202S); anti-phospho-p70 S6 kinase (1:1000 Cell Signalling Technologies #9234); anti-4EBP1 (1:1000 Cell Signalling Technologies #9644); anti-phospho-4EBP1 T70 (1:1000 Cell Signalling Technologies #9455T); anti-SIRT1 (1:1000 Proteintech #60303-1-IG); anti-PGC1α (1:1000 Proteintech #66369-1-IG); anti-NRF1 (1:1000 Proteintech #66832-1-Ig); anti-TFAM (1:1000 Proteintech #22586-1-AP); anti-MT-ND1 (1:1000 ABClonal #A5250).

On the next day, membranes were incubated for 1 h in the dark at RT with the appropriate HRP-conjugated secondary antibodies: anti-rabbit HRP conjugated secondary (1:5000 Cell Signalling Technologies Cat. N°7074S) or anti-mouse HRP conjugated secondary (1:5000 Cell Signalling Technologies Cat. N°7076S). Proteins were detected with an ECL System from Amersham. All western blot raw images are provided in Supplementary Materials.

### Proliferation

#### Sulforhodamine B (SRB) staining

7-10 × 10^3^ cells/well were seeded in 96-well plates and after 24 h the respective media containing the specific treatments were changed. Plates were incubated at 37 °C in humidified atmosphere with 5 % CO_2_ for 24 h, 48 h, 72 h or 96 h. At the given time-point, cells were washed once with PBS solution and fixed with 10 % TCA for 1 h at 4 °C. Then, TCA solution was removed, wells were washed three times with distilled water and after air drying, plates were stained with 0.2 % SRB in 1 % acetic acid solution for 15 min at room temperature. Then, the SRB solution was removed, wells were washed three-times with 1 % acetic acid and after air drying, 100 µL of Tris-base pH 10.4 solution was added in each well for solubilization under shaking during 20-30 min. Plates were read at 490 nm in a spectrophotometer Multiskan KC (Thermo Scientific).

#### Crystal Violet staining

7-10 × 10^3^ cells/well were seeded in 96-well plates and after 24 h the respective media containing the specific treatments were changed. Plates were incubated at 37 °C in humidified atmosphere with 5 % CO_2_ for 24 h, 48 h or 72 h. At the given time-point, cells were washed once with PBS solution and fixed with 4 % formaldehyde for 15 min at room temperature. After letting cells air dry completely, wells were stained with 0.05 % crystal violet solution for 10 min at room temperature and then washed three-times with distilled water. After air drying, the staining was solubilized with 10 % acetic acid and placed in a shaker for 20-30 min. Plates were read at 595 nm in a spectrophotometer Multiskan KC (Thermo Scientific).

#### Clonogenic assay

Approximately five hundred cells were seeded in 60 cm petri dishes cultured in full cell culture medium. The cells were allowed to grow for about 2-3 weeks and medium was changed every three-days. After this period, cells were washed, fixed with 4 % paraformaldehyde, and further stained with 0.05 % crystal violet solution. The plates were scanned for counting the number of colonies by using Image J software (NIH).

#### Anchorage-independent colony formation

Six-well plates were coated with 1 mL of 1.6 % low melting agarose (Sigma-Aldrich) in 2x concentrated cell culture medium and stored at 4 °C for at least 30 min to let the agarose solidify. Afterwards, 1 × 10^5^ cells/well were suspended in 1 mL of 0.8 % low melting agarose medium mixture and seeded. Plates were then incubated at 4 °C for 15 min to allow solidifying the upper layer of the gel. Complete cell culture medium was added on top of the wells and plates were incubated at 37 °C in humidified atmosphere of 5 % CO_2_ for 2–3 weeks. Number of colonies were counted using Image J software.

#### Microspheroids formation

Melanoma cells were resuspended in complete cell culture medium and 3 × 10^4^ cells/well were seeded in ninety-six U shaped well plates (Corning Costar 3799) pre-coated with 1 % agarose to favour microspheroid formation. Cells were incubated at 37 °C in humidified atmosphere of 5 % CO_2_ for 24 h, 48 h or 72 h and microspheroids were photographed using a EVOS XL Core (Invitrogen). Circularity index was quantified by measuring the microspheroids area using Image J software.

#### Metabolite Extraction and Liquid Chromatography-Mass Spectrometry (LC-MS) for Steady-State Metabolomics Analysis

To quantify relative steady-state levels of intracellular metabolites, cells were extracted and analyzed by LC-MS/MS. Cell lines were incubated for 24 h in their specific cell culture media with or without supplementation with 5 µM CRO15 or DMSO. For siRNA against MELK, cells were treated during 24 h with 50 nM siMELK or siControl. Two different siRNA sequences were used for each condition. Then, media was changed, cells were incubated in fresh media and metabolites were extracted after 24 h.

On the given experimental time, metabolites were extracted on dry ice using 4 mL of pre-chilled 80 % methanol (−80 °C), as previously described^35^. Extracts were clarified by centrifugation at 3,000 x g for 5 min to pellet insoluble debris. The remaining pellet was then re-extracted twice with 0.5 mL of 80 % methanol, each followed by centrifugation at 20,000 x g for 5 min. Supernatants from all three extractions were pooled (total ∼5 mL) and dried under a stream of nitrogen using an N-EVAP evaporator (Organomation, Inc, Associates). Dried extracts were reconstituted in 50 % acetonitrile, vortexed for 30 sec, and centrifuged at 20,000 x g for 30 min at 4 °C. The cleared supernatant was transferred to LC-MS vials for analysis.

Metabolites were separated by hydrophilic interaction chromatography (HILIC) on an Ultimate 3000 HPLC system (Thermo Fisher Scientific) equipped with a Waters XBridge Amide column (2.3 x 100 mm, 3.5 μm) and detected on a Q Exactive high-resolution mass spectrometer (Thermo Fisher Scientific) operated with an electrospray ionization (ESI) source. Mobile phase A consisted of 95 % water/5 % acetonitrile supplemented with 10 mM ammonium acetate and 10 mM ammonium hydroxide (pH 9.0), and mobile phase B was 10 % acetonitrile. The gradient was: 0 min, 15 % A; 2.5 min, 30 % A; 7 min, 43 % A; 16 min, 62 % A; 16.1-18 min, 75 % A; 18-25 min, 15 % A. The flow rate was 150 μL/min. Source parameters were set to: capillary temperature 275 °C, sheath gas 35 (arb units), auxiliary gas 5 (arb units), and spray voltage 4.0 kV. Data were acquired in positive/negative polarity switching mode over an m/z range of 60-900, with full-scan MS1 collected at 70,000 resolution (AGC target 1 x 106; maximum injection time 200 msec). The top five precursor ions were selected for data-dependent MS2 fragmentation using HCD (30 % normalized collision energy) at 17,500 resolution. Metabolites were assigned based on accurate mass and confirmed by comparison to authentic standards (retention time) and/or MS2 fragmentation patterns. Acquisition and quantification were performed using Xcalibur 4.1 and TraceFinder 4.1 (Thermo Fisher Scientific), respectively.

For data analysis, the peak areas previously identified were uploaded in csv file format in MetaboAnalyst 6.0 (https://www.metaboanalyst.ca/ Accessed on 17.12.2025). Values were normalized by med and range scaled. Univariate analysis including FC analysis and t tests was performed within the software. Enrichment pathway analysis was also performed with MetaboAnalyst 6.0 using the Kyoto Encyclopedia of Genes and Genomes (KEGG) database.

#### Transmission Electron Microscopy

Melanoma cells were seeded in 12-well culture plates. At approximately 80 % of confluency, cells were washed twice in PBS and then fixed with 1.6 % glutaraldehyde in phosphate buffer (0.1 M pH 7.4) for 1 h at room temperature. Then cells were rinsed in cacodylate buffer (0.1 M pH 7.4) and post-fixed in 1 % osmium tetroxide (reduced with 1 % potassium ferrocyanide). After being rinsed in distilled water, cells were gradually dehydrated in ethanol, embedded in epoxy resin, and incubated at 60 °C overnight for polymerization. Ultrathin sections (80 nm) were assembled on copper grids and contrasted with uranyl acetate (4 % in water) followed by lead citrate. Sections were observed under a JEOL JEM 1400 electron microscope equipped with a Morada SIS camera. Mitochondrial mass was assessed and quantified using ImageJ software (NIH) and data is represented in sum of total area of mitochondrial branching.

### Respirometry

#### High-Resolution Respirometry

High-resolution respirometry was measured using Clark-type oxygen electrodes (Oroboros O2k, Oroboros Instruments, Austria) at 37 °C in a 2 mL close-chamber system under 750 rpm constant stirrer speed as described previously^36^. Briefly, tissue homogenates of tumor xenografts biopsies were prepared as follows:

Tissue biopsies were thawed on ice and placed on top of a petri dish with ice-cold relaxing and biopsy preservation solution (BIOPS) containing 50 mM K^+^-MES, 20 mM taurine, 0.5 mM dithiothreitol, 6.56 mM MgCl_2_, 5.77 mM ATP, 15 mM phosphocreatine, 20 mM imidazole pH 7.1, 10 mM Ca-EGTA buffer (2.77 mM CaK_2_EGTA + 7.23 mM K_2_EGTA; 0.1 μM free calcium). Then, tissue embedded in BIOPS was cut with a scalpel into small pieces, weighted in a precision scale, and 10-15 mg of tissue were subsequently placed onto a tube containing 1 mL room temperature mitochondrial respiration medium (MiR05) containing 110 mM sucrose, 60 mM K^+^-lactobionate, 0.5 mM EGTA, 3 mM MgCl_2_, 20 mM taurine, 10 mM KH_2_PO_4_, 20 mM HEPES, pH 7.1. Then, the mixture of tissue plus medium was placed into a tissue grinder and quickly grinded until completely homogenised with 1-sec pulses maceration. Tissue homogenates were placed into the pre-calibrated O2k-chamber already containing MiR05 up to a 2 mL final volume per chamber.

Mitochondrial respiratory capacities from biopsies based on oxygen concentration and oxygen consumption were evaluated upon titration of a series of substrates, uncoupler and inhibitors (SUIT protocols, Oroboros Instruments) using the reference protocols 1 and 2 (SUIT-001 and SUIT-002)^36^ at minimum two experimental replicates per sample. Respiratory capacities were expressed as O_2_ flow [amol*s^−1^*mg^-1^ protein] and corrected for residual oxygen consumption. All drugs were purchased from Sigma-Aldrich. DatLab 7 software (Oroboros Instruments) was used for real-time data acquisition at a recording interval of 2 sec and for data analysis.

#### Oxygen Consumption Rate (OCR)

OCR was measured using a Seahorse XF96 extracellular flux analyser (Agilent) in intact cells. Cells were seeded at an initial concentration of 1 × 10^4^ cells/well of a XF96 plate and allowed to adhere for 24 h. A measurement plate containing calibrant solution (100 μL per well) was placed in a CO_2_-free incubator at 37 °C overnight. The next day, this plate was run as calibration. Meanwhile, culture media were removed, cells washed twice with PBS and 100 μL of fresh media was added containing 5 mM glucose, 1 mM pyruvate and 4 mM glutamine. To eliminate CO_2_ residue in the medium, cells were incubated for 1 h *prior* to the experiment at 37 °C with CO_2_ in a non-humidified incubator. After calibration, OCR was assayed by sequential addition of mitochondrial inhibitors and uncoupler. Following stabilization of basal respiration, ATP-synthase inhibitor oligomycin (1 µM, port A) was initially added to evaluate the oxygen consumption independent of mitochondrial ATP production. Then, the protonophore carbonyl cyanide m-chlorophenyl hydrazone (CCCP) was added, forcing the transport of H^+^ throughout the mitochondrial inner-membrane and reaching the maximum mitochondrial respiration (500 nM, port B and C). Finally, 0.5 μM rotenone (inhibitor of CI) plus antimycin A (inhibitor of CIII) were added (port D) to block mitochondrial respiration. At least 6-technical replicates were done in the same plate for each sample and 6-measurements were carried out at baseline and after each injection. Three independent experiments were performed. The OCR value was normalized to cell numbers *per* well.

#### Enzymatic activities

For enzyme activities, 1.0 × 10^7^ cells were harvested using trypsin-EDTA 0.25 %, washed with phosphate saline buffer, and a cell pellet was obtained by centrifugation at 300 g for 5 min and snapped frozen at −80 °C. After collection of all samples, the cell pellets were thawed on ice, suspended in protein extraction buffer containing 20 mM Tris-HCl pH 7.2, 250 mM saccarose, 40 mM KCl, 2 mM EGTA, 1 mg/mL BSA and mechanical cell lysis was performed using a glass potter at 4 °C. After centrifugation at 500 g for 20 min, collection of supernatants, and another centrifugation at 650 g for 20 min, cell homogenates were obtained, and protein concentration was determined using the BCA assay (Thermo Fisher). The activity of mitochondrial Complex I and citrate synthase (CS) were measured using standardized protocols, as described previously^37^. The assays were carried out spectrophotometrically at 30 °C. Activities were measured using 40 µg total protein per sample and expressed in nmol*min^−1^ *mg^−1^ protein.

#### Dehydrogenase activity

Cell dehydrogenase activity was determined using a WST-1 assay (Roche), which is based on cleaving the tetrazolium salt WST-1 to formazan. Briefly, cells were incubated with the WST-1 reagent (1:10) for 1 h and then absorbance was measured on a spectrophotometer Multiskan KC (Thermo Scientific) at 490 nm.

#### Statistics

Statistical significance was determined by unpaired Student’s t-test comparing two independent groups. Multiple groups comparison was performed using ordinary one-way or two-way Anova with Tukey’s or Mann-Whitney post-corrections. Statistical significance was considered at α ≥ 0.05. All experiments were performed with a minimum of three biological replicates, and whenever possible, two or more technical replicates. Prism 9 (GraphPad Software; USA) was used for all statistical calculations.

## ACKNOWLEDGEMENTS

The authors would like to thank the platforms at C3M: Microscopy (Marie Irondelle), Cytometry (Frederick Larbret), and Animal facility (Veronique Corcelle). This research was supported by the INSERM, University Cote D’Azur, Fondation pour la Recherche Medicale (EQU202003010248), INCA PLBIO (2020-114), ANR-23-CE14-0023 (TRANSMET) and Canceropole PACA. Sant’Anna-Silva ACB is a recipient of a postdoctoral fellowship from « Bourses d’Excellence Jeunes Chercheurs 2024 » from the University Cote d’Azur.

## AUTHOR CONTRIBUTIONS

A.C.B.S.-S. and S.R. conceived the project and together with M.C. wrote the manuscript. A.C.B.S.-S. designed, performed and analysed all the experiments, P.A. helped with technical support for the *in vivo* experiments and assisted with the data analysis; S.P., N.D.A., F.B. and F.L. provided technical assistance and scientific discussion; Cy.R. provided CRO15; I.B-S. and M.D.T. performed LC-MS/MS metabolomics analyses and scientific input; H.M. and C.R. provided patients samples; R.R., M.C. and S.R. contributed to the experimental design, data analysis and provided scientific discussion; S.R. and T.P. obtained funding and S.R. supervised the project. All authors read, edited, and approved the manuscript.

## REFERENCES

1. Morandi, A. & Indraccolo, S. Linking metabolic reprogramming to therapy resistance in cancer. Biochimica et Biophysica Acta - Reviews on Cancer at 10.1016/j.bbcan.2016.12.004 (2017).

2. Parmenter, T. J. et al. Response of BRAF-mutant melanoma to BRAF inhibition is mediated by a network of transcriptional regulators of glycolysis. Cancer Discov. (2014) doi:10.1158/2159-8290.CD-13-0440.

3. Hernandez-Davies, J. E. et al. Vemurafenib resistance reprograms melanoma cells towards glutamine dependence. J. Transl. Med. (2015) doi:10.1186/s12967-015-0581-2.

4. Baenke, F. et al. Resistance to BRAF inhibitors induces glutamine dependency in melanoma cells. Mol. Oncol. (2016) doi:10.1016/j.molonc.2015.08.003.

5. Shah, R., Singh, S. J., Eddy, K., Filipp, F. V. & Chen, S. Concurrent targeting of glutaminolysis and metabotropic glutamate receptor 1 (GRM1) reduces glutamate bioavailability in GRM1 þ melanoma. Cancer Res. (2019) doi:10.1158/0008-5472.CAN-18-1500.

6. Chang, C. H. et al. Metabolic Competition in the Tumor Microenvironment Is a Driver of Cancer Progression. Cell (2015) doi:10.1016/j.cell.2015.08.016.

7. Vazquez, F. et al. PGC1α Expression Defines a Subset of Human Melanoma Tumors with Increased Mitochondrial Capacity and Resistance to Oxidative Stress. Cancer Cell (2013) doi:10.1016/j.ccr.2012.11.020.

8. Harel, M. et al. Proteomics of Melanoma Response to Immunotherapy Reveals Mitochondrial Dependence. Cell (2019) doi:10.1016/j.cell.2019.08.012.

9. Janostiak, R. et al. MELK Promotes Melanoma Growth by Stimulating the NF-κB Pathway. Cell Rep. (2017) doi:10.1016/j.celrep.2017.11.033.

10. Xu, Q. et al. MELK promotes Endometrial carcinoma progression via activating mTOR signaling pathway. EBioMedicine (2020) doi:10.1016/j.ebiom.2019.102609.

11. Zhang, Z. et al. Upregulated MELK Leads to Doxorubicin Chemoresistance and M2 Macrophage Polarization via the miR-34a/JAK2/STAT3 Pathway in Uterine Leiomyosarcoma. Front. Oncol. (2020) doi:10.3389/fonc.2020.00453.

12. Ren, L., Guo, J. si, Li, Y. heng, Dong, G. & Li, X. yang. Structural classification of MELK inhibitors and prospects for the treatment of tumor resistance: A review. Biomedicine and Pharmacotherapy at 10.1016/j.biopha.2022.113965 (2022).

13. Jaune, E. et al. Discovery of a new molecule inducing melanoma cell death: dual AMPK/MELK targeting for novel melanoma therapies. Cell Death Dis. (2021) doi:10.1038/s41419-020-03344-6.

14. Rambow, F. et al. Toward Minimal Residual Disease-Directed Therapy in Melanoma. Cell (2018) doi:10.1016/j.cell.2018.06.025.

15. Beke, L. et al. MELK-T1, a small-molecule inhibitor of protein kinase MELK, decreases DNA-damage tolerance in proliferating cancer cells. Biosci. Rep. (2015) doi:10.1042/BSR20150194.

16. Zhang, G. et al. Targeting mitochondrial biogenesis to overcome drug resistance to MAPK inhibitors. J. Clin. Invest. (2016) doi:10.1172/JCI82661.

17. Barbato, A. et al. Integrated genomics identifies MiR-181/TFAM pathway as a critical driver of drug resistance in melanoma. Int. J. Mol. Sci. (2021) doi:10.3390/ijms22041801.

18. Long, G. V. et al. Combined BRAF and MEK Inhibition versus BRAF Inhibition Alone in Melanoma. N. Engl. J. Med. (2014) doi:10.1056/nejmoa1406037.

19. Moreno, B. H., Parisi, G., Robert, L. & Ribas, A. Anti-PD-1 Therapy in Melanoma. Seminars in Oncology at 10.1053/j.seminoncol.2015.02.008 (2015).

20. Gray, D. et al. Maternal embryonic leucine zipper kinase/murine protein serine-threonine kinase 38 is a promising therapeutic target for multiple cancers. Cancer Res. (2005) doi:10.1158/0008-5472.CAN-04-4531.

21. Nakano, I. et al. Maternal embryonic leucine zipper kinase is a key regulator of the proliferation of malignant brain tumors, including brain tumor stem cells. J. Neurosci. Res. (2008) doi:10.1002/jnr.21471.

22. Wang, Y. et al. MELK is an oncogenic kinase essential for mitotic progression in basal-like breast cancer cells. Elife (2014) doi:10.7554/eLife.01763.

23. Badouel, C., Chartrain, I., Blot, J. & Tassan, J. P. Maternal embryonic leucine zipper kinase is stabilized in mitosis by phosphorylation and is partially degraded upon mitotic exit. Exp. Cell Res. (2010) doi:10.1016/j.yexcr.2010.04.019.

24. Kuner, R. et al. The maternal embryonic leucine zipper kinase (MELK) is upregulated in high-grade prostate cancer. J. Mol. Med. (2013) doi:10.1007/s00109-012-0949-1.

25. Haq, R. et al. Oncogenic BRAF regulates oxidative metabolism via PGC1α and MITF. Cancer Cell (2013) doi:10.1016/j.ccr.2013.02.003.

26. Laplante, M. & Sabatini, D. M. MTOR signaling in growth control and disease. Cell at 10.1016/j.cell.2012.03.017 (2012).

27. Saxton, R. A. & Sabatini, D. M. mTOR Signaling in Growth, Metabolism, and Disease. Cell at 10.1016/j.cell.2017.02.004 (2017).

28. Sun, C. et al. Reversible and adaptive resistance to BRAF(V600E) inhibition in melanoma. Nature (2014) doi:10.1038/nature13121.

29. Hugo, W. et al. Genomic and Transcriptomic Features of Response to Anti-PD-1 Therapy in Metastatic Melanoma. Cell (2016) doi:10.1016/j.cell.2016.02.065.

30. Herzig, S. & Shaw, R. J. AMPK: Guardian of metabolism and mitochondrial homeostasis. Nature Reviews Molecular Cell Biology at 10.1038/nrm.2017.95 (2018).

31. Leone, R. D. et al. Glutamine blockade induces divergent metabolic programs to overcome tumor immune evasion. Science (80-.). (2019) doi:10.1126/science.aav2588.

32. Mangalhara, K. C. et al. Manipulating mitochondrial electron flow enhances tumor immunogenicity. Science (80-.). (2023) doi:10.1126/science.abq1053.

33. Talantov, D. et al. Novel genes associated with malignant melanoma but not benign melanocytic lesions. Clin. Cancer Res. (2005) doi:10.1158/1078-0432.CCR-05-0683.

34. Jerby-Arnon, L. et al. A Cancer Cell Program Promotes T Cell Exclusion and Resistance to Checkpoint Blockade. Cell (2018) doi:10.1016/j.cell.2018.09.006.

35. Yuan, M. et al. Ex vivo and in vivo stable isotope labelling of central carbon metabolism and related pathways with analysis by LC–MS/MS. Nat. Protoc. (2019) doi:10.1038/s41596-018-0102-x.

36. Doerrier, C. et al. High-resolution fluorespirometry and oxphos protocols for human cells, permeabilized fibers from small biopsies of muscle, and isolated mitochondria. in Methods in Molecular Biology (2018). doi:10.1007/978-1-4939-7831-1_3.

37. Spinazzi, M., Casarin, A., Pertegato, V., Salviati, L. & Angelini, C. Assessment of mitochondrial respiratory chain enzymatic activities on tissues and cultured cells. Nat. Protoc. (2012) doi:10.1038/nprot.2012.058.

