## Supplementary Figure 1 for "MELK controls tumor metabolism to promote resistance to melanoma therapy"

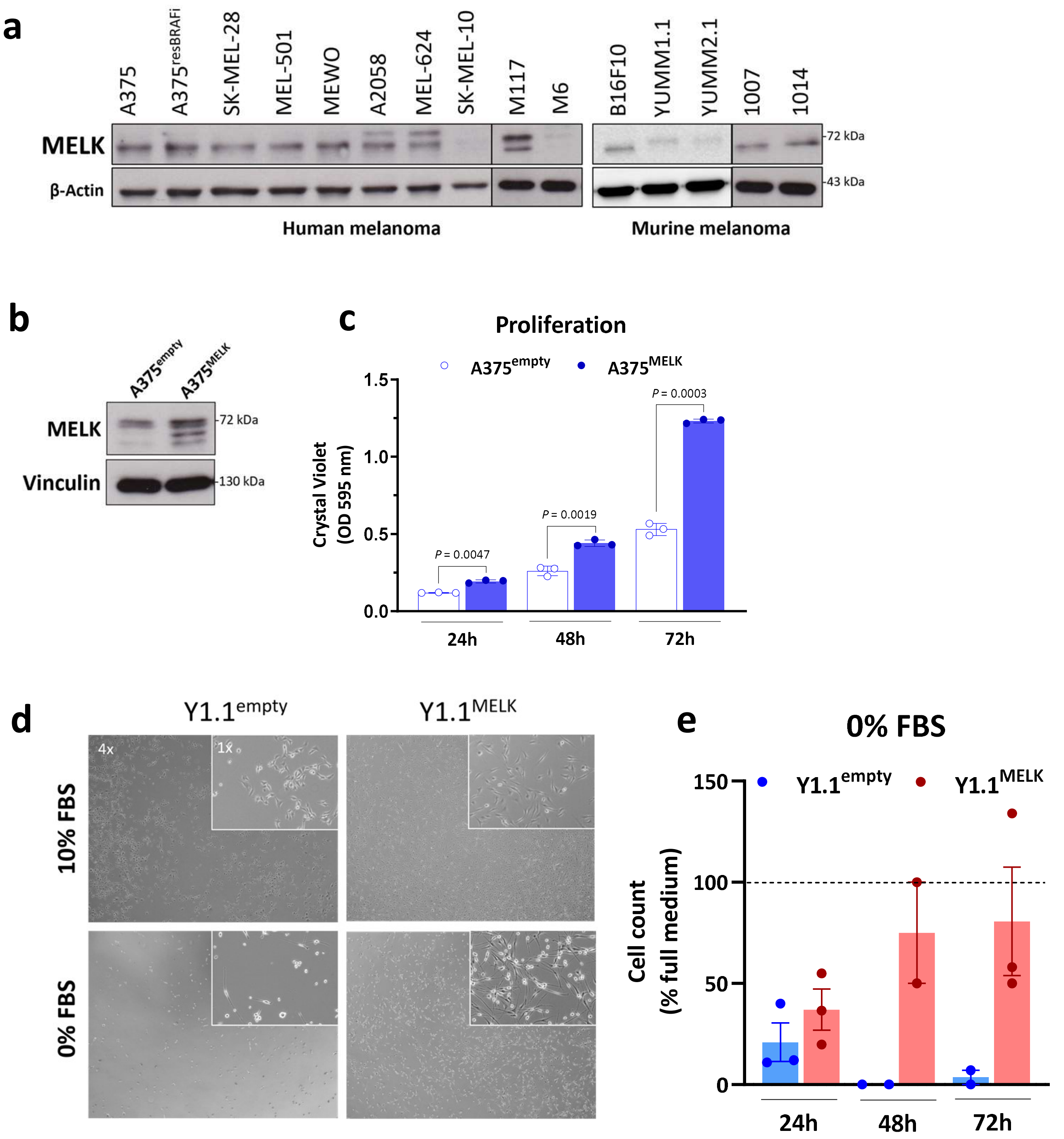


**SUPPLEMENTARY FIGURE 1: (a)** Immunoblot showing MELK expression across murine and human melanoma cell lines; **(b)** MELK overexpression in A375 melanoma cells; **(c)** Proliferation assay with A375 overexpressing MELK or empty vector over time; **(d)** Microscopy pictures of Y1.1 empty vector or MELK cells at basal culture conditions under 10 % FBS or at starvation at 0 % FBS; **(e)** Relative cell count of cells kept at 0 % FBS overtime. Bars indicate ± SEM; N ≥ 3; *P* values and biological replicates are indicated in the graphs.
