## Supplementary Figure 2 for "MELK controls tumor metabolism to promote resistance to melanoma therapy"

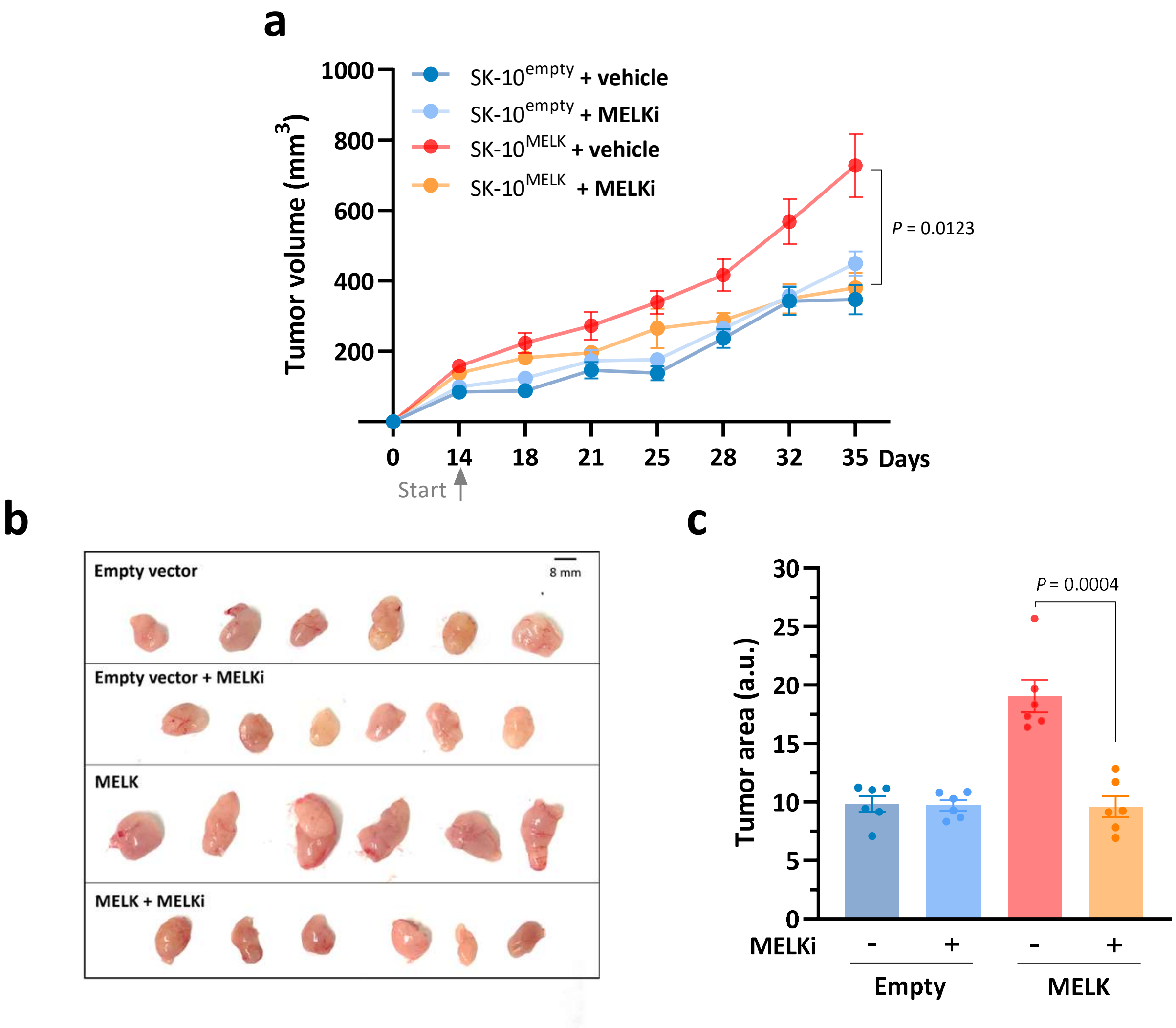


**SUPPLEMENTARY FIGURE 2:** Immunodeficient BALB/c nude mice were inoculated subcutaneously with $5 \times 10^6$ SK-MEL-10 overexpressing MELK or empty control and treated with MELK inhibitor CRO15 (35 mg/kg/day) or vehicle daily. **(a)** Tumor growth curves were followed by measuring the tumor volume ($mm^3$); **(b)** Tumor sizes, and **(c)** *Ex vivo* measurements of tumor areas. Bars indicate ± SEM; N ≥ 3; *P* values and biological replicates are indicated in the graph.
