## Supplementary Figure 3 for "MELK controls tumor metabolism to promote resistance to melanoma therapy"

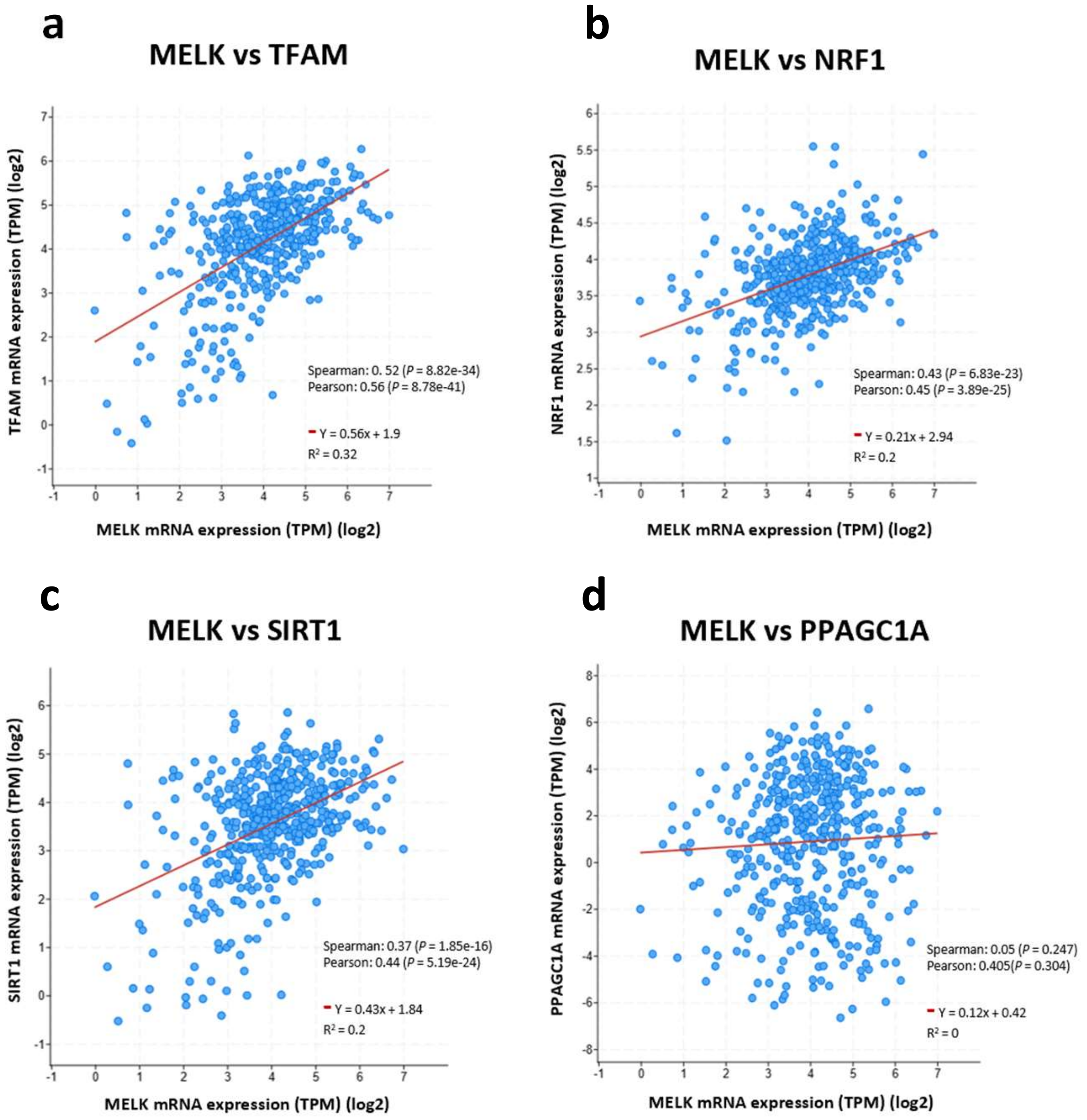


**SUPPLEMENTARY FIGURE 3:** TCGA data base analysis using SKCM cohort showing MELK mRNA expression correlated with the mitochondrial biogenesis markers **(a)** TFAM; **(b)** NRF1; **(c)** SIRT1; **(d)** PPAGC1A. Data are represented in log2 scale; Spearman and Pearson correlation values are displayed in the graphs with respective *P* values.
