## Supplementary Figure 4 for "MELK controls tumor metabolism to promote resistance to melanoma therapy"

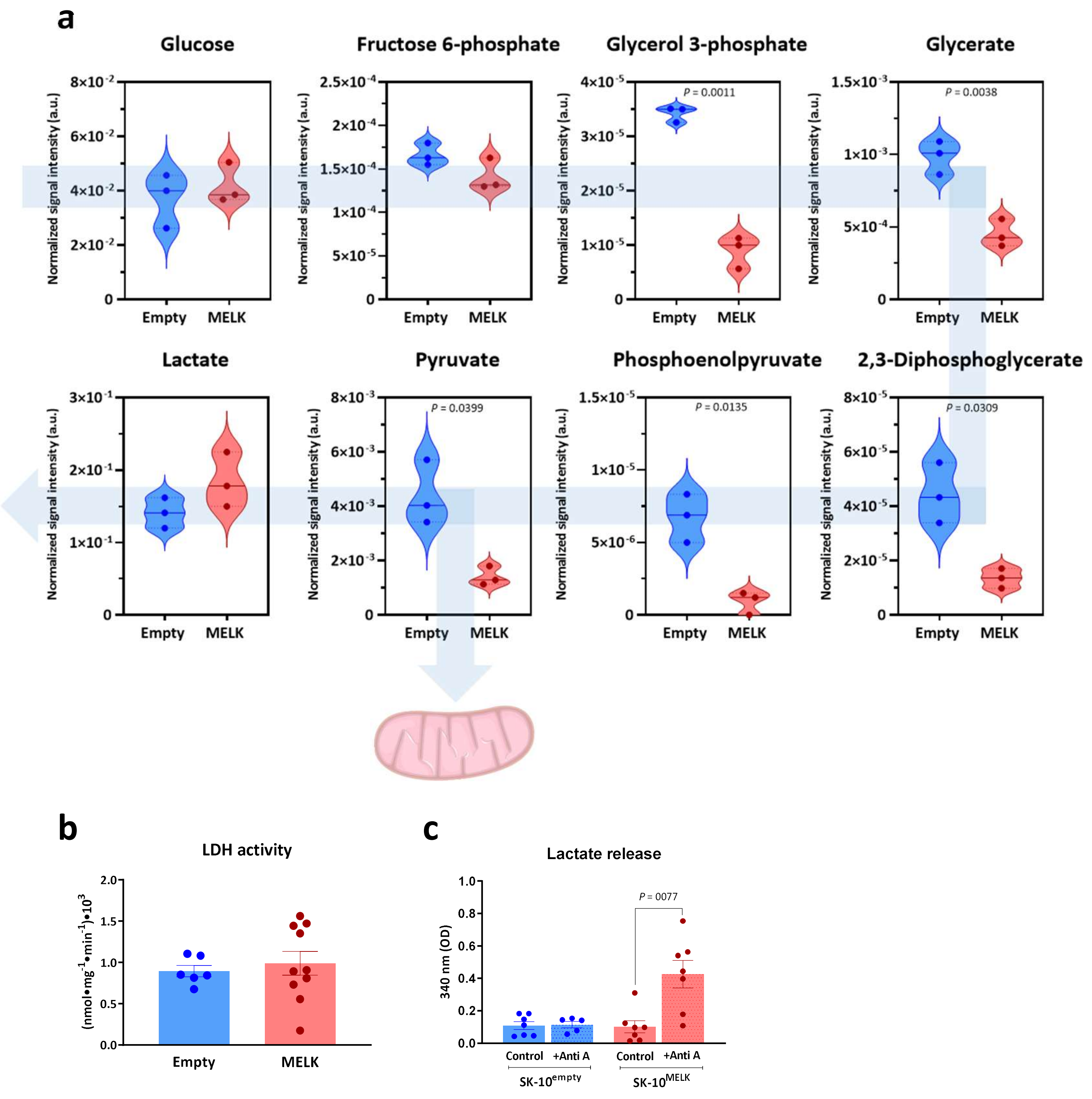


**SUPPLEMENTARY FIGURE 4: (a)** Normalized signal intensity of untargeted steady-state metabolomics analysis displaying glycolytic intermediates concentration comparing SK-MEL-10 expressing empty vector (Empty) and MELK cells; **(b)** Extracellular lactate release of SK-MEL-10 MELK or empty vector cells with or without antimycin A treatment; **(c)** Lactate dehydrogenase activity between SK-10 empty vector or MELK cells (nmol*mg$^{-1}$*min$^{-1}$). Bars indicate ± SEM; N ≥ 3; *P* values and biological replicates are indicated in the graphs. Image provided by Servier Medical Art (https://smart.servier.com/), licensed under CC BY 4.0 (https://creativecommons.org/licenses/by/4.0/).
